# DIB-BOT: An open-source hardware approach for high throughput droplet interface bilayer deposition

**DOI:** 10.1101/2024.01.26.577347

**Authors:** Alexander F Mason, Shelley FJ Wickham, Matthew AB Baker

## Abstract

Droplet interface bilayers (DIBs) provide a controlled lipid environment for the single-molecule investigation of a range of biologically relevant membrane-bound processes and have garnered attention for their potential applications in bottom-up artificial cells, biosensing, and biophysics. However, the fabrication of DIBs is currently hindered by time-consuming processes and specialized equipment. These fabrication limitations prevent the scale-up of DIB assays, making it difficult to generate the large data sets required to achieve statistically significant conclusions in single-molecule biological assays where heterogeneous behaviour is often observed. This research describes an open-source solution, dubbed “DIB-BOT,” constructed by coupling a nanoinjector with an entry-level 3D printer. We present DIB-BOT as a platform to achieve rapid, reproducible, and reliable fabrication of large numbers of DIBs, addressing the limitations of manual methods. Leveraging commercially available off-the-shelf components, DIB-BOT exhibits high spatial reproducibility, minimal user input, and the ability to scale experiments rapidly. Here we demonstrate the utility of the system by integrating pairwise droplet assembly with a fluorescence plate-reader to execute a biologically relevant assay. When compared with manual DIB fabrication, the DIB-BOT had a tenfold reduction in droplet volume error, a threefold reduction in positional error, and 100% droplet yield. Overall, this method has potential to reduce entry barriers to the use of DIB methods, broadening the applications of DIB research, and generating higher quality data sets.

## Introduction

One goal in the field of bottom-up artificial cells is to recapitulate fundamental biological processes in synthetic model systems.^1^ In such models, we can incrementally tune the level of complexity by the stepwise addition of abiotic components towards a “minimal cell”, and along the way design increasingly lifelike biophysical models to answer fundamental questions in biology.^2^ The cell membrane is a fundamental element of such model systems. There are many approaches to forming synthetic membrane mimics in artificial cell platforms, using materials such as polymers, protein bioconjugates, and phospholipids.^3^ In particular, droplet interface bilayers (DIBs) offer an attractive method with unique advantages such as generating planar lipid bilayers with relatively high kinetic stability,^4^ and the ease with which physiologically-relevant asymmetric bilayers can be assembled.^5^

DIBs are formed by touching together two water droplets immersed in a solution of lipid molecules in oil (typically hexadecane).^6–8^ The contact between lipid monolayers surrounding each droplet results in the spontaneous formation of a lipid bilayer at the interface between the two droplets. This technique allows the creation of stable, cell-sized lipid bilayers without the need for a solid support or complicated fabrication methods involving microfluidics.^9^ DIBs have been shown to have similar electrochemical properties^10,11^ to biological membranes and are capable of reconfiguration^12,13^ and ion transport^12^ making them ideal for use in biosensing and drug discovery applications. However, the main limitations of DIBs is that their fabrication can be a time-consuming process that requires specialized equipment and expertise.^14^

To date, there have been limited efforts to increase the throughput and reduce the assembly time for arrays of droplet bilayers.^15,16^ Bilayers are typically assembled via pairwise interaction of droplets where a bilayer is assembled between two droplets. Thus, to increase throughput automation of the delivery of very small volumes (pL to nL) of fluid in a precise location is required. This has been demonstrated using high-precision x/y-translation stages (with associated high cost),^17^ or the repurposing of 3D printers and integration with high-cost flow control.^18,19^ The main limitation of these approaches are the complexity of assembly, relatively high initial cost, and the difficulty of exchanging the aqueous phase to rapidly explore a large parameter space.

Here, we describe the construction of an open-source experimental platform using commercially available off the shelf components to enable rapid, reproducible, and reliable fabrication of large numbers of DIBs. This was achieved by coupling a nanoinjector with an entry level filament deposition 3D printer, which acts as a cartesian robot with a 0.1 mm resolution in x, y, and z. This device, nicknamed “DIB-BOT”, has high spatial reproducibility, can produce large numbers of DIBs in a short amount of time and with minimal user input, and can produce networks of DIBs through simple software modifications. We demonstrate its capacity to answer biologically relevant research questions by integration with existing plate-reader technology to improve access to and increase uptake of DIB analytical methods.

## Results

The generation of DIBs is a two-step process. First, a suitable amphiphile is dissolved in the oil phase. A common formulation for stable synthetic bilayers1,2-diphytanoyl-3-sn-phosphatidylcholine (DPhPC) dissolved in a linear hydrocarbon such as hexadecane. Next, small volumes of aqueous solution are deposited into the oil phase, and when two (or more) water-in-oil droplets are brought into contact, a bilayer will spontaneously form. It is possible to generate DIBs entirely by hand, using a P2 or P10 air-displacement micropipette set to the lowest setting, dispensing droplets on the order of 0.1 - 0.5 μL (100 – 500 nL).^20^ Deposition of an aqueous droplet in an oil solution from a pipette tip requires the operator to raise the tip up through the oil/air interface, whereby the aqueous droplet will shear off from the pipette into the sample well.

While this method is straightforward and requires negligible investment in new equipment, forming DIBs in this way has three major drawbacks. First, operating at the lower limit of a micropipette introduces significant volume variations, and secondly the action of drawing the pipette up through the oil/air interface by hand sometimes results in the generation of more than one droplet. Finally, it is difficult to control the spatial positioning of droplets by hand. Examples of this are shown in Figure 1D, where a micropipette was used to deposit single 0.1 μL aqueous droplets containing 50 μM fluorescein in 16 wells of a 384-well plate. A large variation of droplet sizes was observed (∼50%, 199 ± 99 μm), as well as a lack of control over the number of droplets (∼50% error, 2.1 ± 1.1) and their position relative to the centre of the well (>230 μm, Table 1).

**Table 1:** Comparative performance of single droplet deposition between manual and automatic methods. (#) For automatic droplets all 16 wells only had 1 droplet per well. (†) Uncertainty in droplet diameter is 1 s.d.(‡) Variation in position was calculated by first finding the difference between the x/y co-ordinates of each droplet centroid and the mean of all droplet centroids, then calculating 1 s.d. of these values.

|  | Number of droplets/well<sup>#</sup> | droplet diameter<sup>†</sup> | Variation in Position<sup>‡</sup> |
| --- | --- | --- | --- |
| **Manual (by hand)** | $2.1 \pm 1.1$ | $199 \pm 99 \mu\text{m}$ | X: 235 $\mu\text{m}$<br>Y: 239 $\mu\text{m}$ |
| **Automatic (DIB-BOT)** | 1<sup>#</sup> | $273 \pm 7 \mu\text{m}$ | X: 83 $\mu\text{m}$<br>Y: 62 $\mu\text{m}$ |

**Figure 1:**
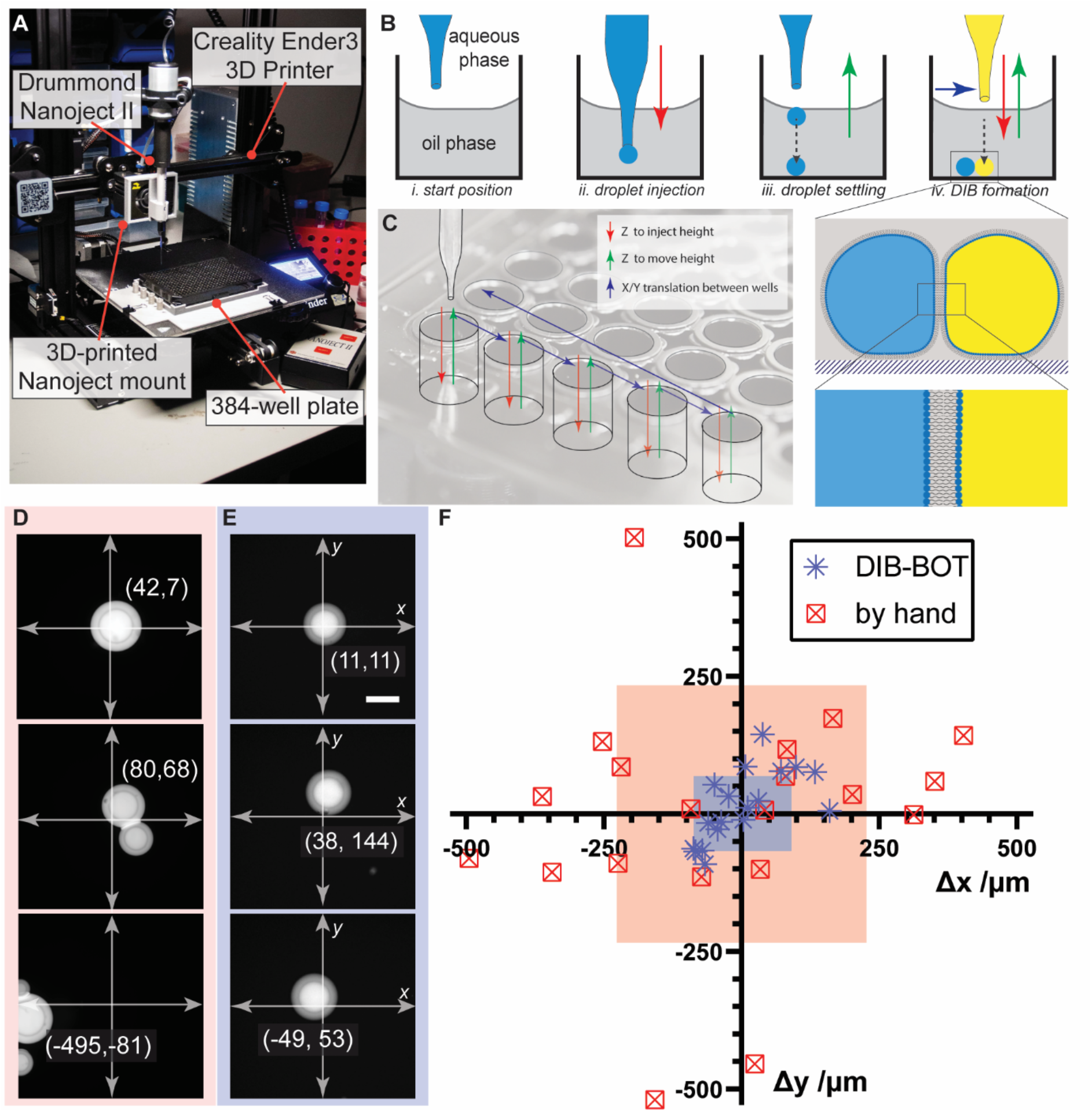
**A**. A photo of DIB-BOT labelled with the key components. **B**. A 2D side-view illustration of the droplet formation process. At the start of every droplet deposition cycle, the microcapillary filled with the aqueous phase starts above the oil/air interface (i). In the next step, the microcapillary is lowered into the oil phase, and the NanojectII is instructed to inject, causing a hanging droplet to be pushed out (ii). In the final step, this hanging droplet is sheared off the microcapillary by withdrawing it upwards back through the air/oil interface (iii). The water droplet then settles to the bottom of the well as it is negatively buoyant. By repeating this process with the same or different aqueous phase in the microcapillary, a DIB can be formed (iv). **C**. This process can be used to rapidly deposit a large number of droplets by inserting an x/y translate step between each cycle. Representative fluorescence images of droplets deposited by hand (**D**.) or using the DIB-BOT €. Scale bar represents 200 μm. Coordinates indicated are raw centroid values. **F**. Spatial repeatability of droplets deposited using the DIB-BOT compared to droplets deposited by hand using an Eppendorf pipette. Each coordinate is calculated by taking the difference between the centroid and the centre of the sample well, with 1 standard deviation illustrated by the coloured box.

The DIB-BOT was developed to rectify these drawbacks, combining readily available commercial off the shelf components to create a machine capable of reliably depositing single aqueous droplets of a specified volume with a high degree of spatial repeatability. Even entry-level FDM 3D printers utilising wheels in V-slots for linear actuation in the x and y axes are capable of 0.1 mm spatial resolution. In order to deliver the aqueous phase, a Drummond NanojectII was mounted to the x-axis carriage using a custom, 3D-printed harness (Figure 1A). A minor modification was made to the Ender3 that enables communication between the 3D-printer and the NanojectII, essentially enabling the injection of nanolitre volumes (from 4.6 nL to multiples of 50 nL) when a specific sequence of gcode is read by the Ender3. Details of this code are available on the project github (<u>AFMason/DIB-BOT: Your guide to building and using DIB-BOT (github.com)</u>). This, coupled with precise translations in x, y, and z (Figure 1B,C) enabled the precise, reproducible deposition of aqueous droplets in hexadecane solutions of DPhPC. The DIB-BOT was used to deposit 100 nL droplets in a 384 well plate and achieved an increase in spatial repeatability compared to depositing droplets by hand (Figure 1D-F). Use of the DIB-BOT reduced variability in droplet size by up to 16-fold (∼50% to ∼3%) and in position by 3-fold (x,y avg. 237 μm to 73 μm). Most impressively, the error in droplet number for the DIB-BOT was undetectable for 16 wells, compared to ∼50% error for the manual method. The DIB-BOT has the additional advantages of being semi-automated. Once the aqueous solution is loaded into the NanojectII, DIB-BOT can cycle through all 16 wells in less than 1 minute with no user intervention. This would compare to a total time of at least 5 mins for an experienced manual user.

The DIB-BOT’s ability to deposit single droplets with high repeatability and in a semi-automated process was then extended to two-droplet systems. The high throughput generation of droplet interface bilayers requires two water-in-oil droplets to be touched together to form a bilayer (Figure 1B). DIBs can be formed by hand, either using a handheld micropipette, or by mounting the Nanoject in a micromanipulator. However, this is difficult and slow, as droplets need to be carefully manipulated individually, pushing individual droplets around the sample well until two are touched together. Therefore, a new DIB-BOT method was written to introduce a second aqueous droplet in each well by implementing a rinsing protocol. After the first droplets were deposited in each sample well, the nanoinjector navigated to the side of the 384-well plate, where three 0.5 mL Eppendorf tubes mounted in a custom 3D-printed tube rack were positioned. First, the nanoinjector empties its contents into a waste tube containing hexadecane. Next, the nanoinjector moves to the second tube and fills with MilliQ (or an appropriate buffer solution). It then returns to the waste tube and empties again. This process was repeated a further two times before the nanoinjector navigated to the third tube to be filled with the new sample, which contained the contents of the second droplet. On the second pass through each sample well, the droplet was deposited with a 0.1 mm offset in x (compared to the first droplet), which resulted in the successful generation of DIBs in every well (n = 16, Figure 2). The DIBs were consistent in size (409 ± 25 μm), positioning (± 53 μm), and the formation of a bilayer (16/16 bilayers). Furthermore, the rinsing protocol was effective, as no observable crosstalk between fluorescence channels was observed (Figure 2, inset).

**Figure 2:**
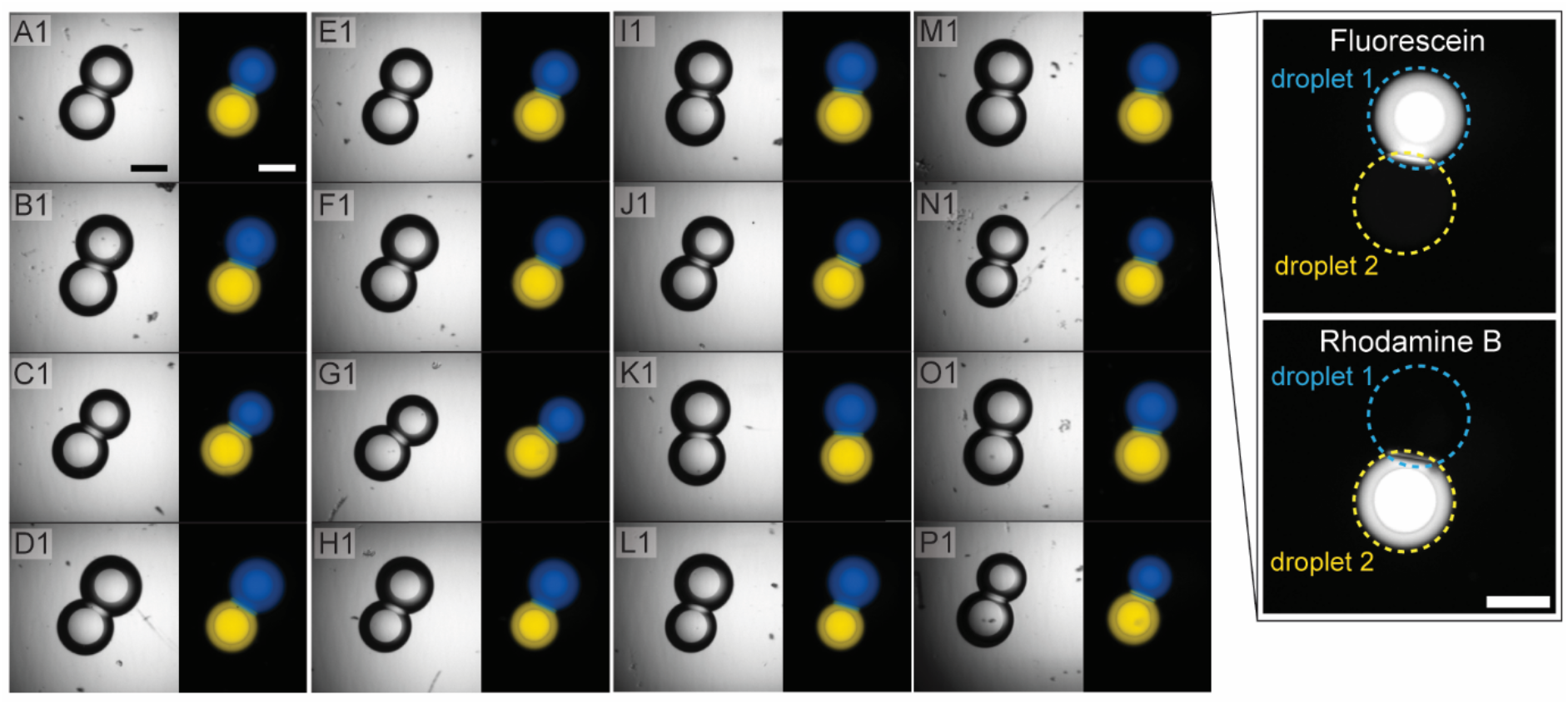
Demonstration of the DIB-BOTs capability to rapidly and consistently generate DIBs in parallel (n = 16), in the discrete sample environments of a 384-well plate. A transmitted light microscopy image is provided for each well, in addition to a merged 2-channel fluorescent micrograph. Blue = droplet 1, 50 μM fluorescein sodium salt, yellow = droplet 2, 30 μM rhodamine B. Scale bar represents 200 μm. **Inset**. Examination of individual fluorescence channels do not reveal any crosstalk between the droplets.

Following the success of the DIB-BOT in reliably generating two-droplet DIBs in a 384-well plate, we tested the capability of the instrument to pattern larger, more complex patterns of DIBs. Patterned multi-DIB systems can be used to model multi-cellular systems, forming synthetic tissues,^21^ but currently such systems are not widely accessible, and do not have the ability to rapidly pattern multiple contents. To this end, a new method was developed where all permutations of three colour droplets were patterned in a 384-well plate, to demonstrate spatial as well as chemical control. The results of this experiment are shown in Figure 3.

**Figure 3:**
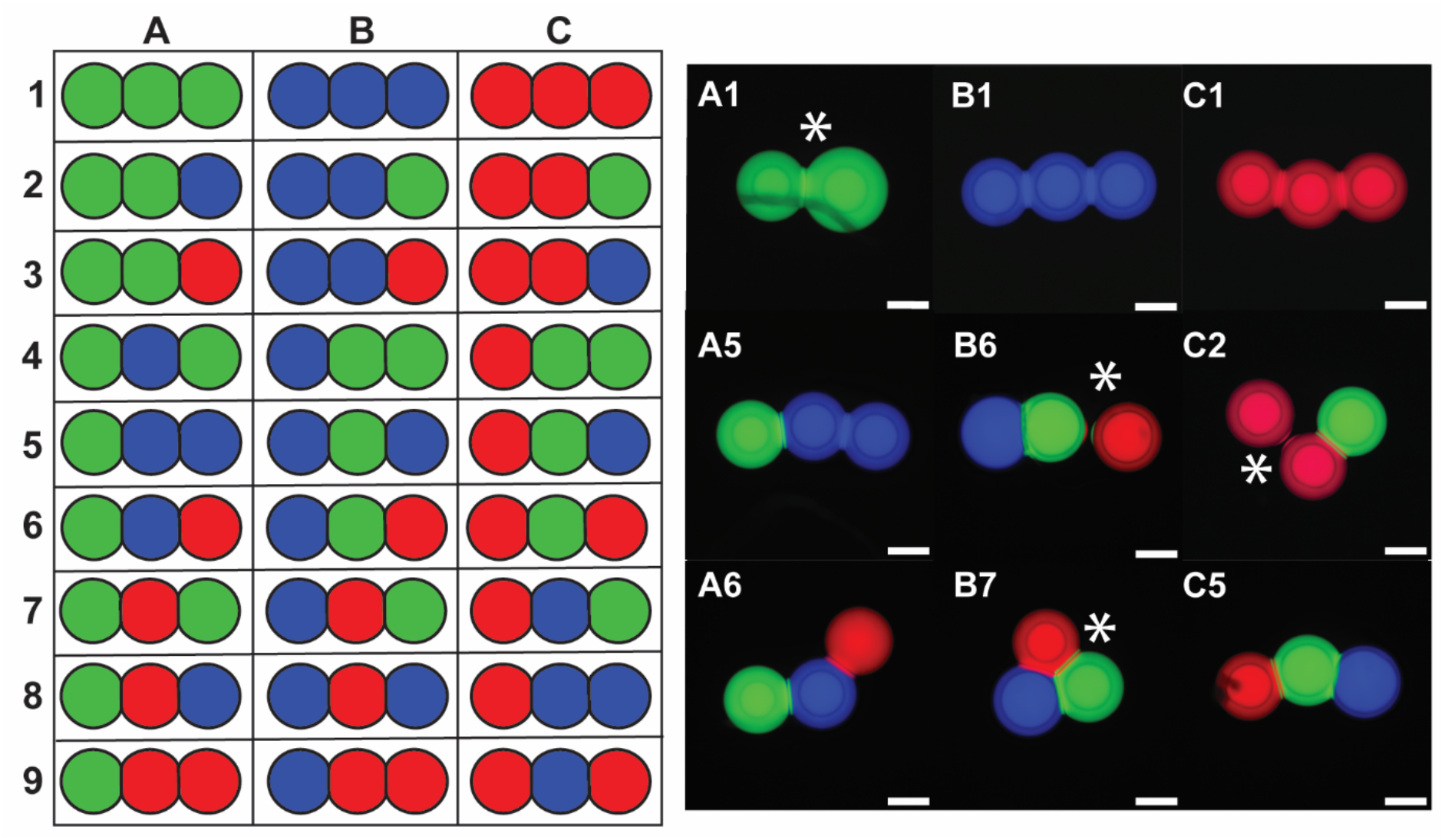
Demonstration of multiplex patterning capability of the DIB-BOT. A representative collection of images for 9 different target patterns is shown. Green = fluorescein, red = Rhodamine B, blue = cy5. Green = 50 μM fluorescein sodium salt, blue = 30 μM rhodamine B, red = 0.5 μM AlexaFluor 647-ssDNA conjugate. Scale bar represents 200 μm. An asterix denotes a droplet network that is functionally incomplete due to absent or incorrect bilayer formation.

Droplet pattern quality was assessed for 3 criteria: number and type of droplets, connectivity of droplets, and geometry. In this case, approx. 30% (8 out of 27) droplet networks could be classified as functionally ‘ideal’ for all 3 criteria, with 3 droplets of the correct type, one bilayer between each droplet, in a line. In many cases (26%) the programmed pattern met the first 2 criteria for type and connectivity of droplets, but varied in geometry such that instead of the designed 180° the line of droplets was instead angled <180°. Such systems still presented the correct order of droplets and thus may still be useful depending on the requirements of the assay. However, incorrect geometry can limit subsequent scaling up to larger droplet arrays, by preventing correct connectivity when further droplets are added. In some cases, the droplets fused prematurely or one of the droplets did not deposit close enough to form a bilayer with its neighbour (44%), making these droplet systems not suitable for use in any assays. Thus, these data show the potential of DIB-BOT to generate large, chemically complex droplet networks in a semi-automated fashion. In principle, this could be expanded to many more colours, as the rinsing and re-loading of the capillary injector is trivial and only takes 1-2 minutes per change.

Finally, we demonstrated the ability of the DIB-BOT to assemble arrays of DIBs that could be used to interrogate the function of a membrane protein. For this, we measured ion flow following the successful insertion of the pore protein α-hemolysin (αHL) which we included in the aqueous solution in the interior of specific droplets. This was achieved by using a slightly modified DIB-BOT method as employed in Figure 2, but with additional nanoinjector rinsing steps to examine multiple experimental conditions in the same run. In this two-droplet experiment, α-hemolysin (αHL, 10 nM) was encapsulated in the first droplet in an aqueous buffer solution containing 0.66 M CaCl_2_. In the second droplet, Fluo-8 (40 μM), a membrane impermeable calcium profluorophore, was encapsulated in buffer containing 1.32 M KCl to preserve osmotic balance across the bilayer. In both negative and positive controls αHL was omitted. For the positive control, 0.66 M CaClwas pre-incubated with Fluo-8 to indicate maximum emission, which was used for subsequent normalisation. Change in Fluo-8 emission, indicating Ca^2+^ flow across the membrane, was measured every 5 minutes using a plate reader (Figure 4). Significant heterogeneity was observed in the results for the same sample in different DIBs. At this concentration of αHL, approximately half of the DIBs showed high activity, where there was a clear increase in Fluo-8 signal indicating pore insertion and Ca^2+^ transport occurred (Fig 4. Red). The other half showed low activity, with stable Fluo-8 signal indicating either pore insertion or transport did not occur (Fig 4. Blue).

**Figure 4:**
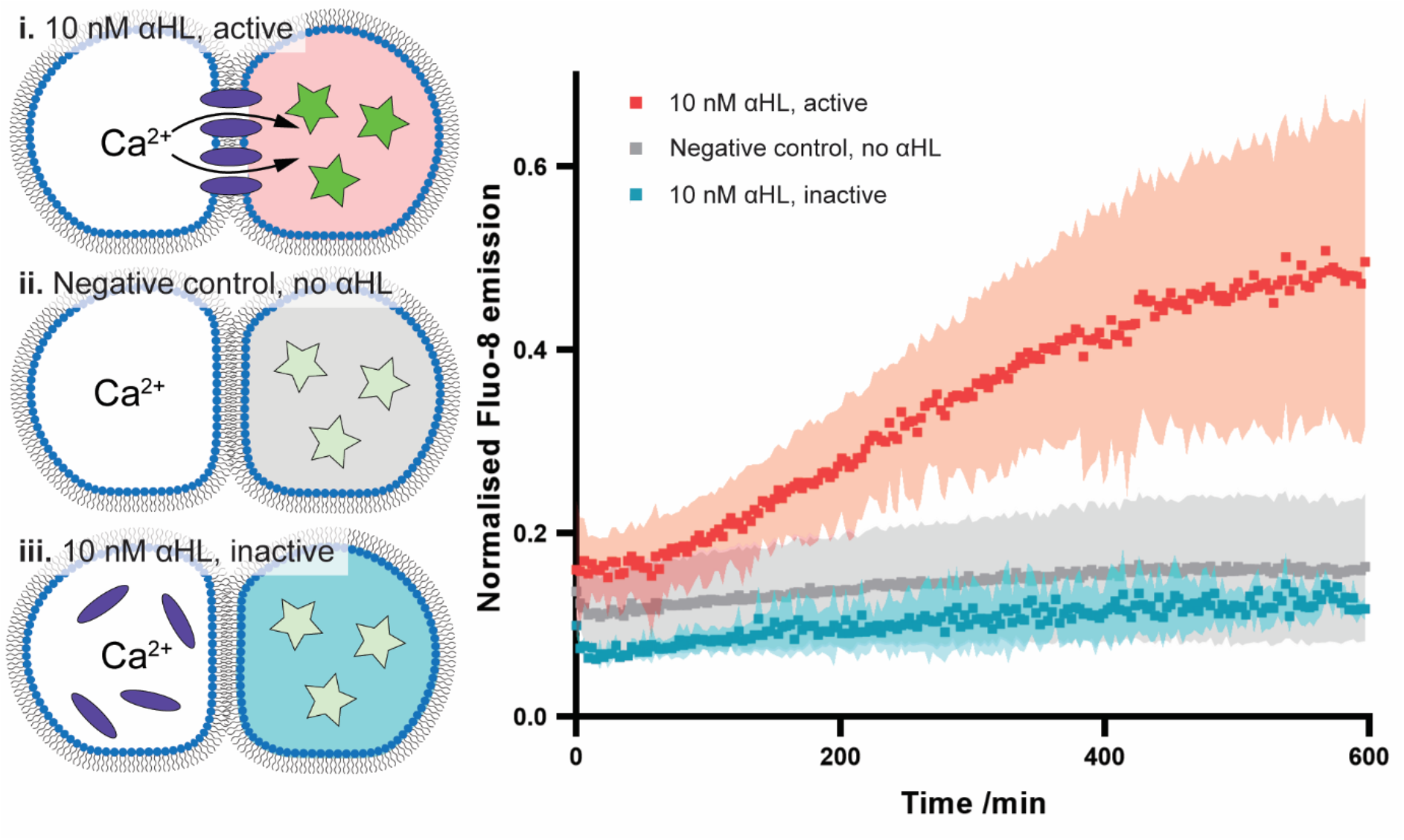
Poration activity of a canonical membrane pore alpha-hemolysin (αHL). The DIB-BOT was employed to generate 16 DIBs containing Ca^2+^ ions in one droplet, separated from a Ca^2+^ fluorophore Fluo-8 in its adjacent droplet. The presence of active αHL enabled the rapid translocation of Ca^2+^(i., n=4), much faster than the negative control with no αHL (ii., n=4). In approximately 50% of αHL droplets, αHL did not show any activity (iii., n=4, separated from αHL sample data set). Emission of Fluo-8 was normalised to a positive control (n=4) in which Ca^2+^ and Fluo-8 were pre-mixed in the same droplet (data not shown).

## Discussion

DIBs can be difficult to generate without access to expensive and/or bespoke equipment. The DIB-BOT is built using relatively low-cost commercial off the shelf components, and all other required components can be printed quickly and cheaply using the 3D printer itself. This lowers the entry cost to utilise synthetic bilayer systems, potentially enabling more researchers around the world to design new experiments utilising DIBs. There are multiple interesting membrane bound processes that would benefit from investigation in a stable environment where lipid composition can be controlled.^4^ Our DIB technology is capable of high-throughput generation of DIBs in discrete chemical environments such as a 384 well plate. This technology could help to elucidate new biophysical phenomena that requires exploring an expansive parameter space, examining factors such as lipid composition, droplet size, protein concentration, substrate/ligand concentration, and droplet network morphology. All these parameters could in principle be explored in an automated or semi-automated fashion with the DIB-BOT.

Nonetheless, there are several current limitations to the DIB-BOT. First, despite being a relatively straightforward build, the construction and operation of DIB-BOT does require some 3D printing, electronics, coding expertise, and expertise in safety evaluation and risk assessment. This may be an obstacle for end users who just want a “turn-key” solution.

Secondly, in membrane protein assays such as in Figure 4, the method suffered from poor reproducibility, with variation between runs for the same experimental conditions. This was likely a consequence of utilising a fluorescence plate reader that did not account for the single point source of signal in each well, as opposed to averaging hundreds or thousands of cells in a typical biological activity assay. Nevertheless, the DIB-BOT shows good potential for the development of high-throughput assays as it was possible to reliably print large numbers of two-colour DIBs.

Thirdly, while the DIB-BOT accurately prints three-droplet DIBs with correct geometry (see Figure 3), it does so with lower yield than its 100% yield for fabricating two-droplet DIBs. This limits the ability to build up larger droplet arrays, where geometrical errors can lead to connectivity errors on addition of subsequent droplets. This is most likely due to the resolution of the x and y axes. The Ender3 is an entry level 3D printer, and while a resolution of 0.1 mm is sufficient for the majority of (simple) 3D printed objects, it does introduce sources of error for the DIB-BOT. One possible workaround for this is to upgrade the linear actuation from roller bearings on V-slots to linear rails. More expensive FDM 3D printers could also provide a better droplet deposition outcome. Another limitation is the fact that only one nanoinjector can be mounted to the 3D printer at a time. This means that any “multi-colour” experiments need to be designed sequentially, with the nanoinjector being rinsed and reloaded between sample changes. This introduces a time delay, makes the process less automatic, and for certain applications a source of error (for example where even a small amount of cross-contamination would be unacceptable). However, the open-source nature of this instruments means that alternate injection apparatus could be installed. The challenge would then be adapting the communication between gcode read by the 3D printer and the injection apparatus.

In this work we have outlined the construction and core capabilities of the DIB-BOT, which extends to the integration of the membrane protein αHL and assessment of its ability to facilitate transmembrane communication in the form of ionic current flow. By increasing throughput of samples, we were able to reveal heterogeneity in protein behaviour in different DIBs and identify 2 distinct populations that were either active or inactive for membrane transport. There is scope to increase the complexity of this experimental setup and work towards more lifelike models, for example by depositing large numbers of droplets to work towards artificial tissues,^17^ tuning the formulation of phospholipids to examine structure-function relationships, and encapsulating in vitro transcription translation (IVTT) systems such as PURExpress® to synthesise proteins of interest in situ.^21,22^ Furthermore, we envisage applications in synthetic biology and biophysics, whereby the DIB-BOT provides an opportunity to rapidly screen membrane proteins such as pores in combination with small molecular activity modulators.

Overall, we have shown that we can clearly distinguish the presence or absence of a pore in a bilayer. This offers promise for low-cost, high-throughput studies of libraries of pores,^23^ ion channels^24,25^ or GPCRs.^26^ For example, in experiments in directed evolution or in protein engineering,^27^ many variants of a pore might be tested for insertion efficacy, or, alternately, a library of membrane proteins could be screened pairwise against a library of ligands, in the other droplet, to determine transmembrane activity. Large libraries could be tested in parallel and then subsequently selected for further optimisation. In the case of DNA-based pores,^28^ these candidate pores are directly made out of DNA and highly amenable to barcoding for subsequent refinement and to test multiple combinations of pore size, activation, and membrane tethering to identify the best route to compatibility between DNA nanostructures and lipid droplet systems.

## Conclusion

Here we have described the development of a semi-automated robot that is capable of reliably generating DIBs in 384-well plates. This has been accomplished using relatively low cost, commercially available components, which has the potential to lower the barriers to entry for researchers looking to design new experiments using DIBs. We have demonstrated a proof-of-concept application of an assay for the evaluation of the membrane protein αHL, paving the way for more rigorous assessment of other more complex membrane bound processes. The current iteration of DIB-BOT is not well suited for the reliable deposition of extended (n≥3) droplet networks, due to limitations in x/y resolution of the Ender3. Nevertheless, the open-source nature of this design enables further modifications and we encourage further refinement of the DIB-BOT.

## Methods

### Materials

Glass capillaries (3-000-203-G, 1.14mm O.D. x 3.5″ length, Drummond) and NanojectII (Drummond) were purchased from Adelab Scientific. Fluo-8, sodium salt (AAT-21088, AAT Bioquest) was purchased from Jomar Life Research. DPhPC, AR-20 silicone oil, CaCl_2_, KCl, HEPES, α-hemolysin and 384-well plates (#781091, 384-well polystyrene flat-well μclear Black, Greiner Bio-One) were purchased from Sigma-Aldrich.

### Modification of the 3D printer

For this project, an Ender 3 printer was selected due to its low cost and open framework (so that the NanojectII would not hit any part of the printer during patterning methods). In principle, any 3D printer with separate x/y/z motion that can facilitate the Nanoject could be used. The gcode method files described herein should be compatible with no (or only minor) changes for most 3D printers.

After the Ender3 was assembled, it was used to 3D print the Nanoject holder (slicing settings and model “Nanoject_holder.stl” can be found on github). After printing, the bridge support and any small printing errors were carefully removed from the Nanoject holder with a scalpel and 120 grit sandpaper.

Next, the motherboard cover was removed to facilitate the installation of the relay that activates the Nanoject II switch on the FAN3 circuit. A detailed explanation of this step and circuit diagram (Figure Sx) can be found in the SI and on the accompanying github (<u>AFMason/DIB-BOT: Your guide to building and using DIB-BOT (github.com)</u>). Before replacing the motherboard cover, run a test method (Inject test.gcode) to see if the connections have been made correctly and that the Ender3 can successfully tell the NanojectII when to inject. During operation of the DIB-BOT, users should wear appropriate PPE, particularly lab safety glasses or goggles to protect eyes from any accidental capillary breakages, or implement an appropriate equivalent safety measure such as a Perspex safety screen.

### Micropipette fabrication

Micropipettes were pulled using a Sutter P-87 capillary puller. The puller was set in the following manner: heat = ramp (672), pull = 0, v = 40, time = 250, pressure = 500. The puller should loop twice, with the capillary breaking on the second loop. If the capillary snaps on the first loop, lower the heat and/or velocity parameters in small increments. Before mounting in the Nanoject, micropipettes were confidently but gently stabbed through a single sheet of kimwipe paper to break the tip cleanly, which was verified using a transmitted light microscope (4x objective).

### Deposition into 384 well plates

Before starting any patterning, it is necessary to 3D print the custom plate holder that assists holding plates in stable position. This ensures that the 384 well plate does not move during the patterning runs, and assists in reproducibility between separate runs.

Powdered DPhPC was dissolved in chloroform to 100 mg/ml, prior to splitting it up into 100 μL aliquots in 2 mL glass HPLC vials. The solvent was removed under a gentle stream of nitrogen to yield a 5 mg film. DPhPC films were further dried in vacuo for at least 18 hours before further use, and stored under nitrogen at −20 °C if not used immediately. Lipid films were re-dissolved in 1:1 hexadecane:AR20 silicone oil to a concentration of 5 mg/mL and sonicated for 30 minutes. Each well to be used in an assay was filled with 13 μL of DPhPC solution.

Inner aqueous solutions were typically prepared at a 10 μL scale in plastic PCR tubes. For the αHL assay in Figure 4, the following solutions were prepared:

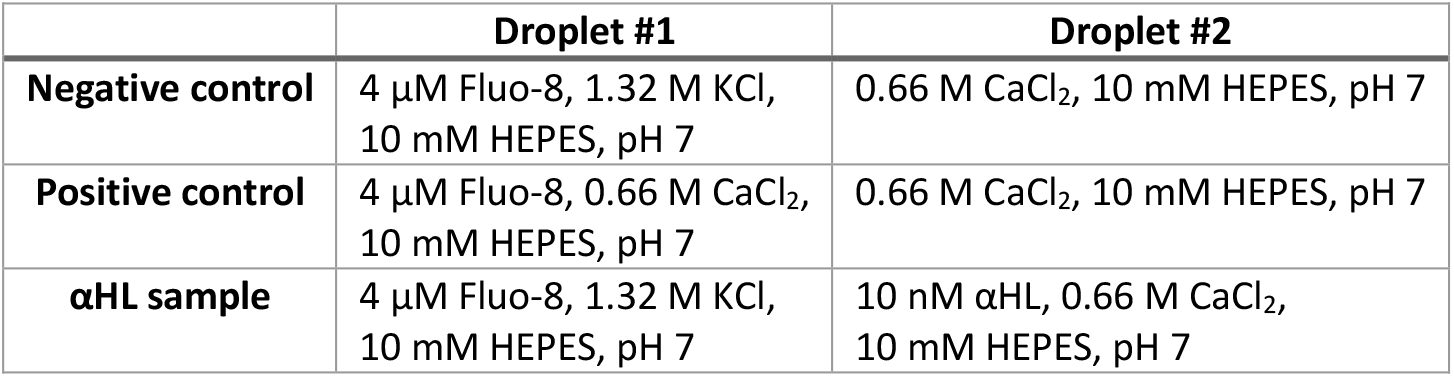

|  | **Droplet #1** | **Droplet #2** |
| --- | --- | --- |
| **Negative control** | 4 μM Fluo-8, 1.32 M KCl, 10 mM HEPES, pH 7 | 0.66 M CaCl<sub>2</sub>, 10 mM HEPES, pH 7 |
| **Positive control** | 4 μM Fluo-8, 0.66 M CaCl<sub>2</sub>, 10 mM HEPES, pH 7 | 0.66 M CaCl<sub>2</sub>, 10 mM HEPES, pH 7 |
| **αHL sample** | 4 μM Fluo-8, 1.32 M KCl, 10 mM HEPES, pH 7 | 10 nM αHL, 0.66 M CaCl<sub>2</sub>, 10 mM HEPES, pH 7 |

The microcapillary was mounted in the NanojectII as per manufacturers instructions, backfilled with 1:1 hexadecane:AR20 silicone oil, and filled with aqueous solution. At pause points in the DIB-BOT methods, the microcapillary was emptied of the previous solution, rinsed 3x with MilliQ, and 1x with the following solution before the next solution was filled. The droplet volume for this experiment was set to 50.6 nL using the binary switches appropriately on the NanojectII control unit.

### Imaging of 384 well plates

A Zeiss Celldiscoverer 7 was used to acquire all droplet images. A Zeiss 5× air objective with 0.5× zoom was utilised, with a binning of 2 × 2 resulting in 2752 × 2208 pixel images. Temperature and COincubation was switched off. For kinetic experiments, images were acquired every 6 minutes for 12 hours, with an image being acquired at the centre of each well, using adaptive focus. In multi-channel experiments, the following settings were used:

Brightfield: 2.2% intensity, 1.9 ms exposure

Fluo-8/Fluorescein: 8.9% power @ 470 nm, 30 ms exposure, 514/30 nm emission filter Rhodamine B: 14.6% power @ 567 nm, 70 ms exposure, 592/25 nm emission filter Cy5: 10% power @ 625 nm, 150 ms exposure, 709/100 nm emission filter

### Plate reader analysis

A BMG Labtech CLARIOstar plate reader was used to measure αHL kinetic curves. The instrument was run in kinetic mode, with a cycle time of 180 seconds over 200 cycles (10 hours total). Excitation filter was set to 483/14 nm, dichroic filter to 502.5 nm, and emission filter to 530/30 nm. Gain was set to 1000 and a focal height of 11 mm was selected. Measurements were run at 25°C with no shaking or mixing.

